# Motif analysis in co-expression networks reveals regulatory elements in plants: The peach as a model

**DOI:** 10.1101/2020.02.28.970137

**Authors:** Najla Ksouri, Jaime A. Castro-Mondragón, Francesc Montardit-Tardà, Jacques van Helden, Bruno Contreras-Moreira, Yolanda Gogorcena

**Author notes:** Senior authors (Bruno Contreras-Moreira, Yolanda Gogorcena).

## Abstract

Identification of functional regulatory elements encoded in plant genomes is a fundamental need to understand gene regulation. While much attention has been given to model species as *Arabidopsis thaliana*, little is known about regulatory motifs in other plant genera. Here, we describe an accurate bottom-up approach using the online workbench RSAT::Plants for a versatile ab-initio motif discovery taking *Prunus persica* as a model. These predictions rely on the construction of a co-expression network to generate modules with similar expression trends and assess the effect of increasing upstream region length on the sensitivity of motif discovery. Applying two discovery algorithms, 18 out of 45 modules were found to be enriched in motifs typical of well-known transcription factor families (bHLH, bZip, BZR, CAMTA, DOF, E2FE, AP2-ERF, Myb-like, NAC, TCP, WRKY) and a novel motif. Our results indicate that small number of input sequences and short promoter length are preferential to minimize the amount of uninformative signals in peach. The spatial distribution of TF binding sites revealed an unbalanced distribution where motifs tend to lie around the transcriptional start site region. The reliability of this approach was also benchmarked in *Arabidopsis thaliana*, where it recovered the expected motifs from promoters of genes containing ChIPseq peaks. Overall, this paper presents a glimpse of the peach regulatory components at genome scale and provides a general protocol that can be applied to many other species. Additionally, a RSAT Docker container was released to facilitate similar analyses on other species or to reproduce our results.

**One sentence summary:** Motifs prediction depends on the promoter size. A proximal promoter region defined as an interval of -500 bp to +200 bp seems to be the adequate stretch to predict *de novo* regulatory motifs in peach

## 1. Introduction

Peach [*Prunus persica* (L.) Batsch], a member of *Prunus* genus, is one of the best genetically characterized species within the Rosaceae family. With a small size diploid genome (2n = 2x =16; 230 Mbp), and relatively short generation time (2-3 years), peach has become a model species for fruit genetic studies (Abbott et al., 2002). Obtaining elite genotypes with broad environmental adaptations and good fruit quality are the fundamental targets of all *Prunus* breeding programs, since they directly affect the economical relevance of this crop (Gogorcena et al., 2020). Indeed, previous works have reported strong affinity between environmental cues and the fruit quality and aroma (Wong et al., 2016; Tanou et al., 2017). To stand the environmental stimuli and ensure edible fruit development, a complex re-arrangement of the gene expression network is required.

The modulation of gene expression is a complex process occurring at various levels from which the transcriptional regulation is the core control code (Petrillo et al., 2014). The transcription machinery is regulated by an interplay between DNA-binding proteins called transcription factors (TFs) and cis-regulatory elements (CREs). TFs bind short sequences known as TF binding sites (TFBS) or motifs located at CREs (e.g., promoters, enhancers, silencers). TFs may act as either activators or repressors of gene expression, leading to dynamic changes of the cellular pathways. For peach, annotation of TFs is available in the plant transcription factor database (plantTFDB) (Tian et al., 2019).

As of February 2020, plantTFDB v5.0 stores 2780 peach TFs classified into 58 families (http://planttfdb.cbi.pku.edu.cn/). While much is known about TF families, TF-binding motifs remain elusive. Deciphering the cis-regulatory network has become a prerequisite toward scoping out the foundations of transcriptional regulation in *P. persica*. The computational exploration of these DNA motifs has been greatly stimulated by the availability of genomic data and the release of whole genome sequence assemblies (Verde et al., 2013; Verde et al., 2017). In this context, a variety of plant motif finders has emerged. Notwithstanding their value, they are hampered by certain limitations such as, a restricted range of species, Promzea for maize (Liseron-Monfils et al., 2013), and AthaMap for *Arabidopsis* (Steffens et al., 2005), and limited analysis capabilities around experimentally defined motifs as PlantCare, (Rombauts et al., 1999) or PlantPAN, (Chang et al., 2008). Thereby, to circumvent these pitfalls, we have adopted a plant-customized tool for *de novo* motifs discovery, RSAT::Plants (http://rsat.eead.csic.es/plants/). RSAT has both a friendly user interface and command-line tools for versatile analyses in a wide collection of plants (Nguyen et al., 2018). Since the analysis of proximal promoter regions is easier in small genomes with short intergenic regions, most of cis-regulatory motif predictions so far have been conducted in *Arabidopsis thaliana* (Ma et al., 2012; Korkuc et al., 2014; Cherenkov et al., 2018).

In *P. persica* there are only two examples of regulatory motif discovery, in particular on a set of 350 dehydrin promoter sequences (Zolotarov and Strömvik, 2015) and 30 heat responsive genes (Gismondi et al., 2020). In contrast to these case studies, we propose a structured bottom-up framework to identify statistically over-represented motifs on a genome scale. Our probabilistic approach relies on the hypothesis that genes within co-expressed modules are likely co-regulated by the same TFs. This approach has been successfully tested in other species, for example in *Arabidopsis thaliana* (Koschmann et al., 2012; Ma et al., 2013) and maize (Yu et al., 2015). According to Bianchi et al., 2015, an arbitrary defined segment of 1500 bp upstream of the transcription start site (TSS) can be considered as the proximal promoter in peach. However, recent studies about the genomic delimitation of proximal promoters in *Prunus persica* effectively reduced this region to a window of approximately 500 nt (Montardit-Tardà, 2018).

The proposed approach relies on three fundaments, i) an accurate definition of co-expressed gene modules, ii) an assessment of the effect of upstream region length regarding the effectiveness of motif discovery and, finally iii) disclosing the effect of splitting the analysis around the TSS site in discovering potential cis-elements. All together, we demonstrate the utility of our strategy in analyzing genome wide data to provide insights on gene regulation dynamics across tissues and specific conditions. To the best of our knowledge, no work has been reported on cis-elements present in *P. persica* on genome wide level, hence the originality of our survey. Additionally, the predicted motifs from this study can be browsed at (https://eead-csic-compbio.github.io/coexpression_motif_discovery/peach/), where we provide readers with direct links to the results, source code and a Docker container to reproduce the analysis on any other plant species.

## 2. Results

### 2.1 Identification of differentially expressed transcripts and construction of weighted co-expression network

After quality assessment and pseudo-alignment, an expression matrix was generated from eight peach published transcriptomes, including treated and control samples with their corresponding biological replicates. Differential analysis yielded 11,335 altered transcripts using *Q*-value < 0.01 and |β| > 1 thresholds. The number of differentially expressed transcripts (DETs) identified in each RNA-seq experiment is listed in **Table 1**. Detailed information about quality control, pseudo-alignment and differential expression analyses is shown in **Table S1**. An overview of our workflow is provided in **Figure 1**.

**Table 1.** Summary of RNA-seq data used as input and the number of differentially expressed transcripts (DETs) identified in each RNA-seq experiment.

| Project ID | References | Experiments | Tissues | Conditions | DETs |
| --- | --- | --- | --- | --- | --- |
| PRJNA271307 | (Li et al., 2015) | Ripening stage | Fruit | 6 | 2601 |
| PRJNA288567 | (Sanhueza et al., 2015) | Cold storage | Fruit | 6 | 6447 |
| PRJNA248711 | (Bakir et al., 2016) | Hyper hydricity | Leaf | 2 | 15 |
| PRJEB12334 | (Ksouri et al., 2016) | Drought | Root/Leaf | 4 | 350 |
| PRJNA252780 | (Jiao et al., 2017) | Low T° | Stigma | 2 | 406 |
| PRJNA323761 | Unpublished | Drought | Root | 2 | 1118 |
| PRJNA328435 | Unpublished | Cold storage | Fruit | 2 | 2963 |
| PRJNA397885 | Unpublished | Chilling injury | Fruit | 4 | 2429 |

**Figure 1.**
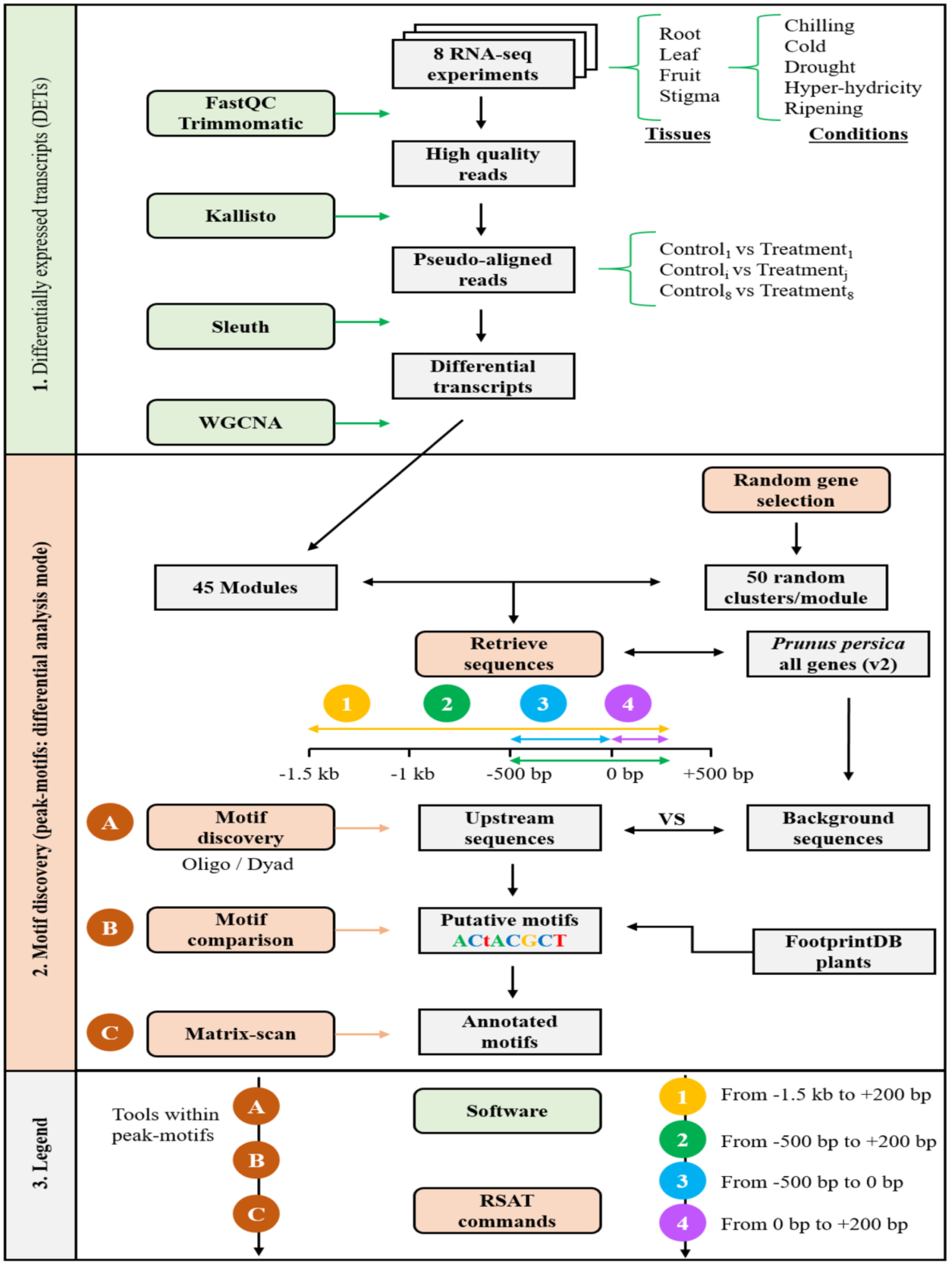
Bottom-up framework for *de novo* motif discovery. **<u>Step1:</u>** differential expression analysis for transcript detection and extraction of co-expressed modules. **<u>Step2</u>:** *de novo* motif detection using the peak-motifs tool from RSAT::Plants. Numbers correspond to the different tested upstream tracts, with TSSs anchored on position 0 bp, while letters represent tools within peak-motifs. Green and orange boxes label software and RSAT tools, respectively.

The WGCNA R-package was adopted to construct an unsigned co-expression network for 11,335 stress-related transcripts. All samples and DETs were considered in the network construction, as neither outliers nor transcripts with missing values, were detected (**Figure S1. A**). Using a dynamic tree cut algorithm, 45 co-expression modules were retained with size ranging from 29 to 1795 transcripts per module (**Figure S1. C)**. The 45 distinct modules (labeled with different colors) are shown in a dendrogram in which major tree branches constitute modules and leaves correspond to DETs (**Figure S1. B**).

### 2.2 Transcription factor binding site (TFBS) prediction

#### 2.2.1 Effect of proximal promoter length on prediction accuracy

As a first step towards extracting regulatory signatures, upstream region boundaries were defined from -1500 bp to +200 bp relative to TSS (Up 1). Six out of 45 modules were found to display positive signals and higher significance when compared to the random clusters. Upstream regions of modules (M9, M10, M11, M18, M21 and M41) matched known core DNA-binding elements corresponding to Myb-like, BZR, CAMTA, bZip, E2FE, and TF families. Modules with their corresponding regulatory elements are represented in **Figure 2** and further information is provided in **Table S2**. Motifs resulting from both oligo and dyad analysis correspond to signatures with strong confidence estimation. Besides, eight poly (AT)-rich signals were discarded from M1, M2, M3, M4 and M6 due to their low complexity. Curiously, these (AT) patterns were also detected in the random clusters and their occurrence seemed to be associated with the size of the module (**Table S3**). For instance, M1 is the largest module with 1795 sequences and (AT)-repetitive signals were detected in 40 out of the corresponding 50 random clusters.

**Figure 2.**
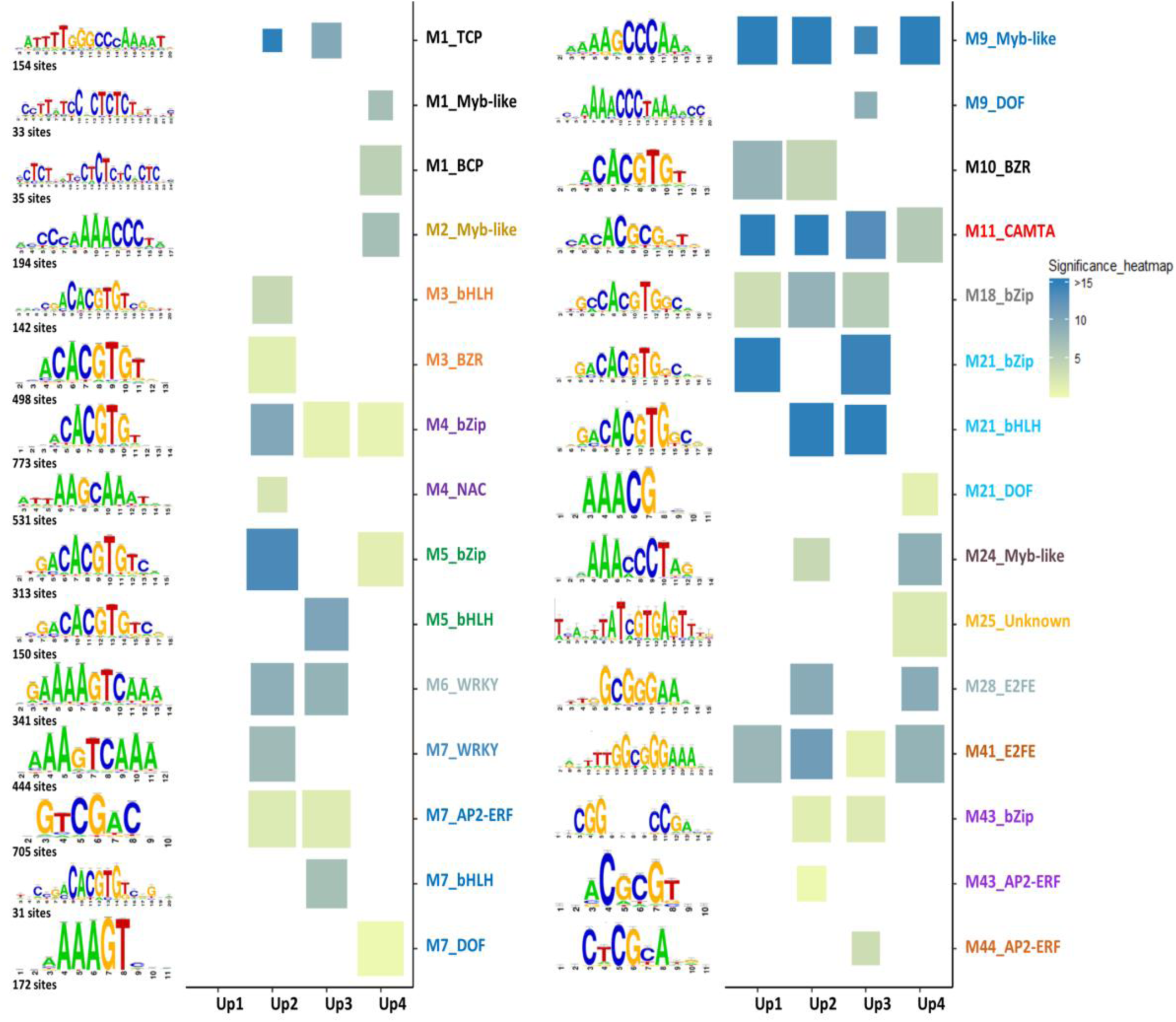
Position Specific Scoring Matrix (PSSM) representation of top scored discovered motifs per modules, along different upstream lengths. The x-axis corresponds to the four intervals: Up 1: [-1500 bp, +200 bp], Up 2: [-500 bp, -200 bp], Up 3: [-500 bp, 0 bp] and Up 4 [0 bp, +200 bp]. The y-axis informs about the motif family revealed per module. Cell colors indicate the statistical significance of the identified motifs while cell sizes represent the normalized correlation (Ncor). Number of sites corresponds to the number of sites used to build the PSSM. When motifs from the same family are identified with both algorithms (oligo and dyad-analysis), or in different upstream tracts (Up 1, Up 2, Up 3 and Up 4), only the most significant one is represented in the heatmap. Further details are provided in **Table S3**. An interactive report with source code is accessible at https://eead-csic-compbio.github.io/coexpression_motif_discovery/peach/

Furthermore, when we restricted the motif discovery to the region with [-500 bp, +200 bp] boundaries (Up 2), fifteen modules were found to discern statistically significant motifs. These were then grouped into 10 TF families as illustrated in **Figure 2** (TCP, bHLH, BZR, bZip, NAC, WRKY, AP2-ERF, Myb-like, CAMTA and E2FE).

An in-depth look at the major changes occurring when trimming the upstream segments to 500 bp resulted in interesting observations, summarized as follows. Spurious (AT) rich events considered as low quality predictions were limited to M2 and were replaced by relevant regulatory elements in M1, M3, M4 and M6 (**Table S2**). Significant signals buried in the long upstream region (Up 1) were inferred in modules M24, M28 and M43 (**Figure 2, Table S2**). Besides, shortening the upstream promoter region size to 500 bp enhances the statistical relevance of the predicted motifs, compared to the negative controls, regardless of the algorithm applied.

Overall, these findings suggest that shortening the upstream region increases the signal-to-noise ratio to detect biologically relevant motifs and, at the same time, reduces the occurrence of low complexity AT-rich motifs. In **Figure 3**, we illustrate a clear showcase of this observation. Indeed, with both oligo and dyad analysis, the statistical significance of motif E2FE found in Module M41 (black bars) has noticeably increased compared to those identified in random clusters (gray bars). Hence, more significant motif discovery was accomplished in the window of [-500 bp, +200 bp].

**Figure 3.**
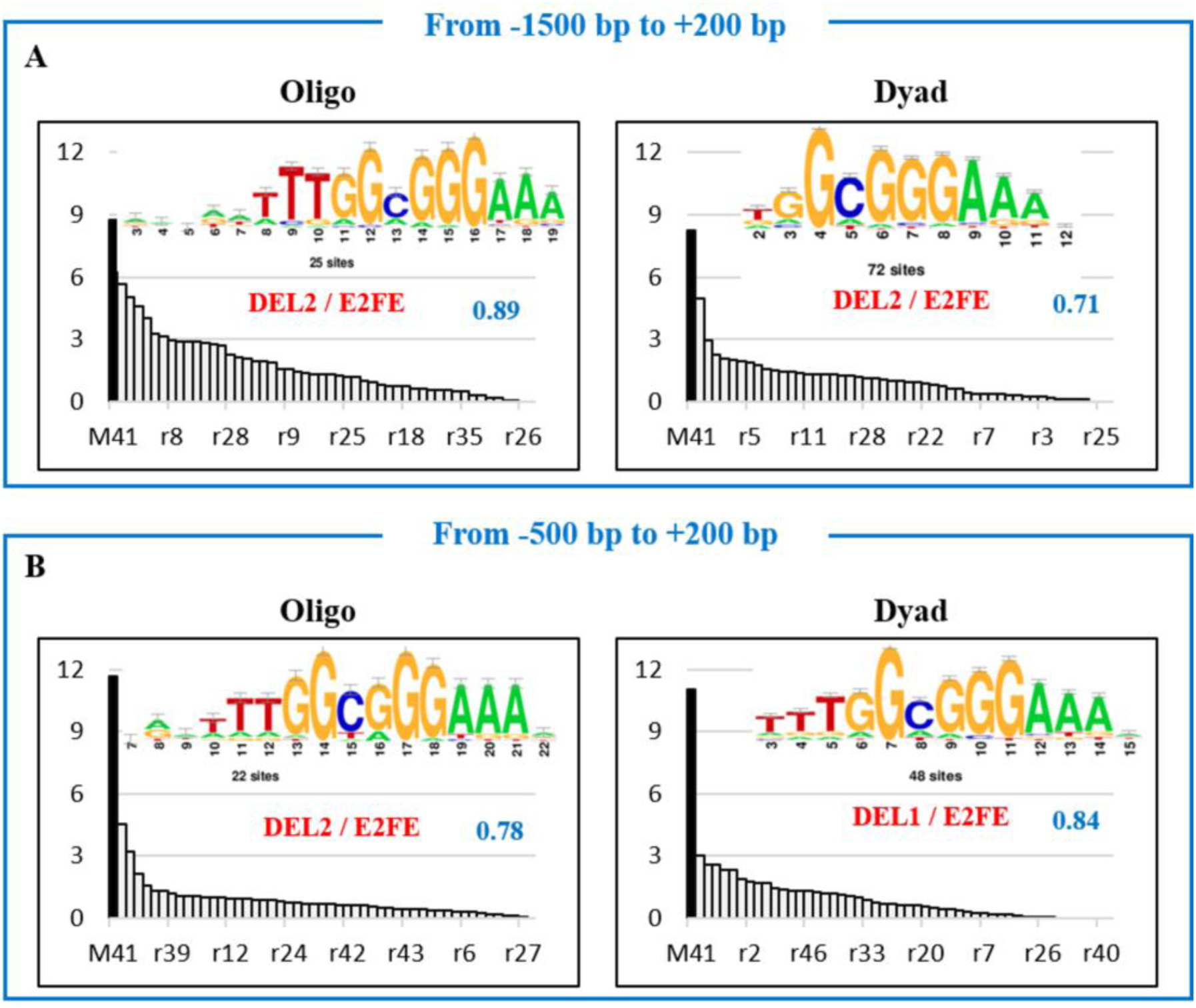
Illustrative comparison between predicted motif DEL2 (corresponding to E2FE transcription factor) within two different upstream promoter lengths: -1500 bp to +200 bp **(A)** and -500 bp to +200 bp **(B)**. The name of the best match among plant motifs in footprintDB is labeled in red, next to its Ncor (Normalized correlation) value labeled in blue. The x-axis corresponds to the module of interest (M41) and random clusters ranked ranked by the most significant motifs. The y-axis corresponds to the statistical significance -log10 (*P*-value). Number of sites corresponds to the occurrence number of a single motif. The evidence supporting the putative motifs is Ncor (in blue) and the significance (black bars) when compared to negative controls (gray bars).

#### 2.2.2 Effect of splitting the promoter region around the TSS on motif prediction

Next, due to the difference in nucleotide composition in coding and non-coding regions, we subdivided the proximal promoter region in two segments around the TSS, with each interval examined separately: upstream, from -500 bp to 0 bp (Up 3), and downstream, from 0 to +200 bp (Up 4). Doing so, motifs of two additional TF families were discovered, BCP in module M1, DOF in modules M7, M9 and M21. In contrast to BCP sites lying downstream the TSS (Up 4), DOF sites were found across both intervals (see **Figure 2, Table S2**). Intriguingly, an uncharacterized motif was over-represented in upstream 4 of module M25 requiring further research.

In conclusion, a total of 77 TF binding motifs were revealed from the different assessed promoter regions (**Table S2**). Modules with candidate predicted motifs might be classified in two types depending on their potentially matching TF. Indeed, across the four examined upstream tracts, we recognize those with motifs bound by a single TF family, considered as single TF-driven modules (e.g., M6, M11, M18, M28 and M41). Conversely, modules having multiple TFBS for several distinct TFs suggest a possible combinatorial regulation under particular circumstances. However, more evidence is needed to address this issue. On the other hand, we observed that the majority of cis-regulatory elements yielded in this study were mainly detected in the upstream region Up 2, defined from -500 bp to + 200 bp (**Figure 2, Table S2)**.

### 2.3 Gene Ontology enrichment

A Gene Ontology analysis was conducted to annotate the potential function of the gene modules. Thirteen modules were significantly enriched with biological processes (**Figure 4**). Six GO terminologies were particularly intriguing and will be briefly described. In modules M1 and M18, transcripts were over-represented respectively in leaf and root tissues under drought experiment which is in line with the “photosynthesis” and “response to water” enrichment. Similarly, module M2 was enriched for “response to stimulus” with high TPM values in fruit tissue at different ripening stage. Transcripts within M5 were mostly abundant in fruit tissue under cold stress, in line with the “cold acclimation” enrichment. Not surprisingly, “response to stress” was over-represented in fruit in module M10 as we are dealing with stress conditions. Finally, hormonal levels are known to imbalance under stress explaining the enrichment of “response to hormone stimulus” in M21. Overall, we consider that the GO enrichment results (**Figure 4.A**) are in harmony with the expression profiles of transcripts in **Figure 4.B**.

**Figure 4.**
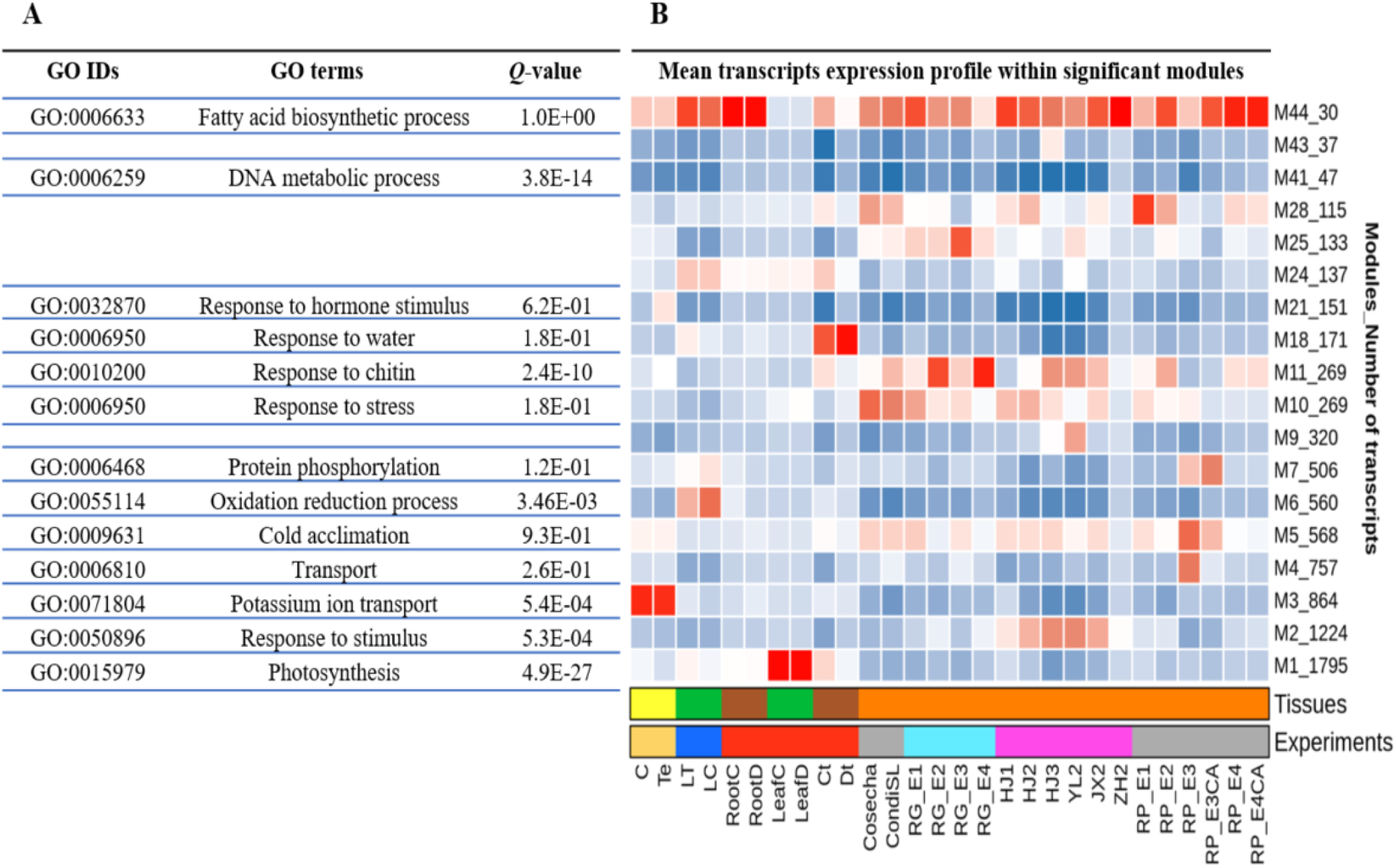
Functional annotation of relevant gene modules. **(A):** Gene ontology enrichment. **(B):** Mean transcript abundance profiling in term of transcripts per million (TPM). The x-axis corresponds the different experimental conditions while the y-axis indicates the number of differentially transcripts per module. Experiment and tissue types are highlighted by different colors (see the color key at the bottom of the figure). Gene profiles along the different conditions are provided at (https://eead-csiccompbio.github.io/coexpression_motif_discovery/peach). See supplementary **Table S1** for the abbreviations.

### 2.4 TFs annotation and prediction of their TFBS using footprintDB

The predicted modules were examined for genes encoding TFs. In total 39 annotated TFs were shortlisted in **Figure 5**. Myb and Myb-like TFs were exclusively expressed in modules M1 and M2. They were particularly over-represented in fruit and leaf tissues in agreement with their transcript profiling illustrated in **Figure 4.B**. We hypothesize that Myb factors may act as regulators of drought stress and ripening in peach. In the same vein, bHLH genes identified in M3 were notably abundant in stigma tissue, which is in accordance with **Figure 4.B**. NAC and E2FE transcription factors were respectively annotated in M4 and M41, and their coding genes were repressed among experiments in all tissues. The WRKY TFs assigned to module M6 were abundant under hyper hydricity fitting with **Figure 4.B** and suggesting a regulatory function of the WRKY in such a condition. Module M7 was associated with genes encoding three TFs with different expression profiles (DOF, bHLH and ERF). Calmodulin binding proteins identified in M11 and bZip annotated in M18 and M21 were highly abundant among all experiments indicating that they may be involved in multiple biological processes.

**Figure 5.**
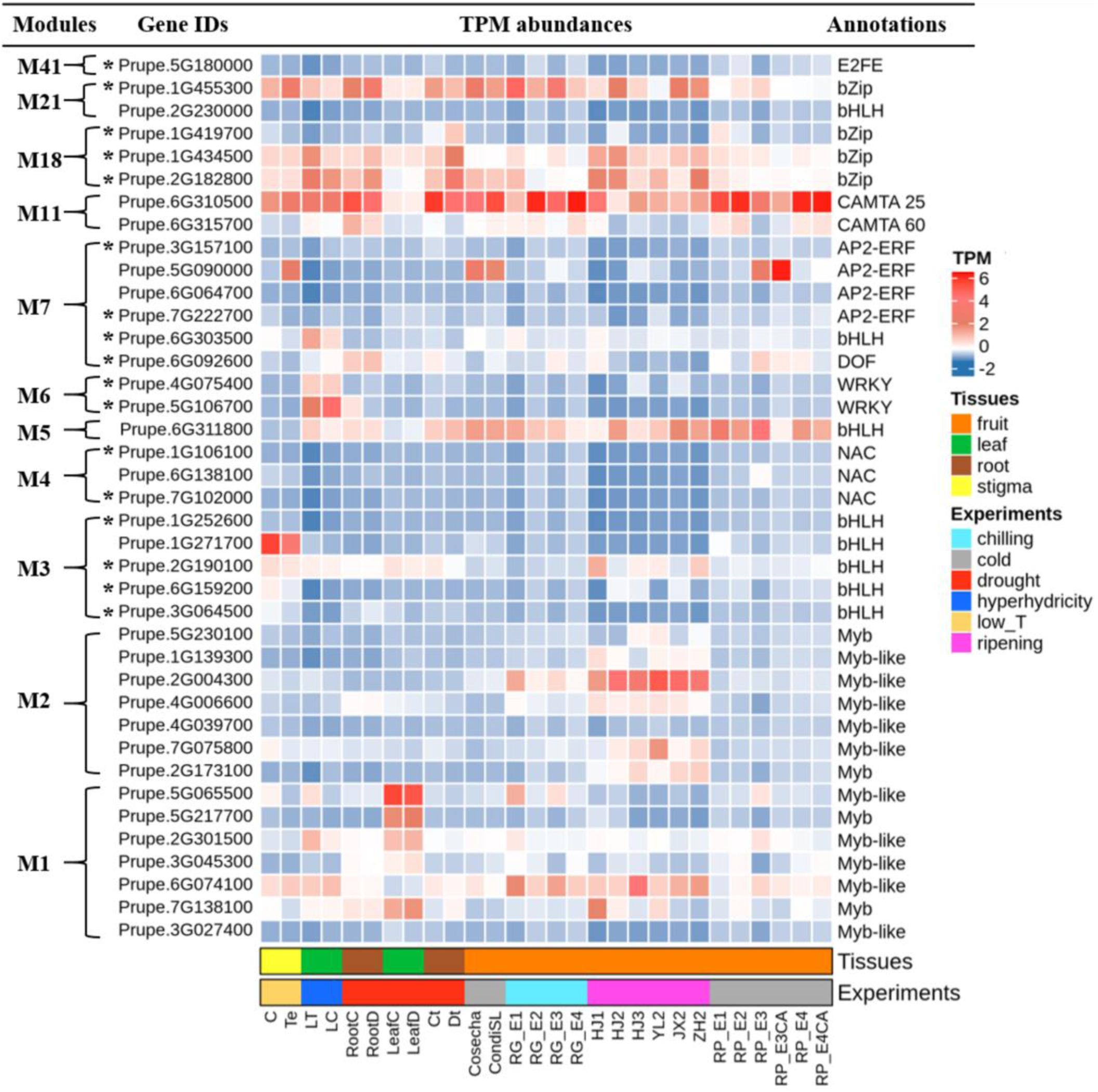
List of transcription factors within relevant modules. Blue and red squares indicate transcripts per million while bottom color bars correspond to the tissues types and different experiments, respectively (See the legend at the right side of the figure). TFs showing sequence similarity between their footprintDB and RSAT predicted motifs are labeled with a star.

Subsequently, we verified whether the disclosed motifs in each module are the actual binding sites of the aforementioned TFs (**Figure 5**). TFs were individually examined for their potential DNA-site using footprintDB and results were compared to those derived from RSAT. Consensus sequences predicted from genes coding TFs showed high similarity to consensus sequences predicted from modules (**Table 2)**. As for instance, the binding motif “tTTGGCGGGAAA” identified in module M41 is almost identical to E2FE-predicted site “TTTTGGCGGGAAAA” from the same module. This suggests that E2FE may modulate gene expression in M41 and “tTTGGCGGGAAA” motif could be the *bona fide* binding site of this transcription factor.

**Table 2.**
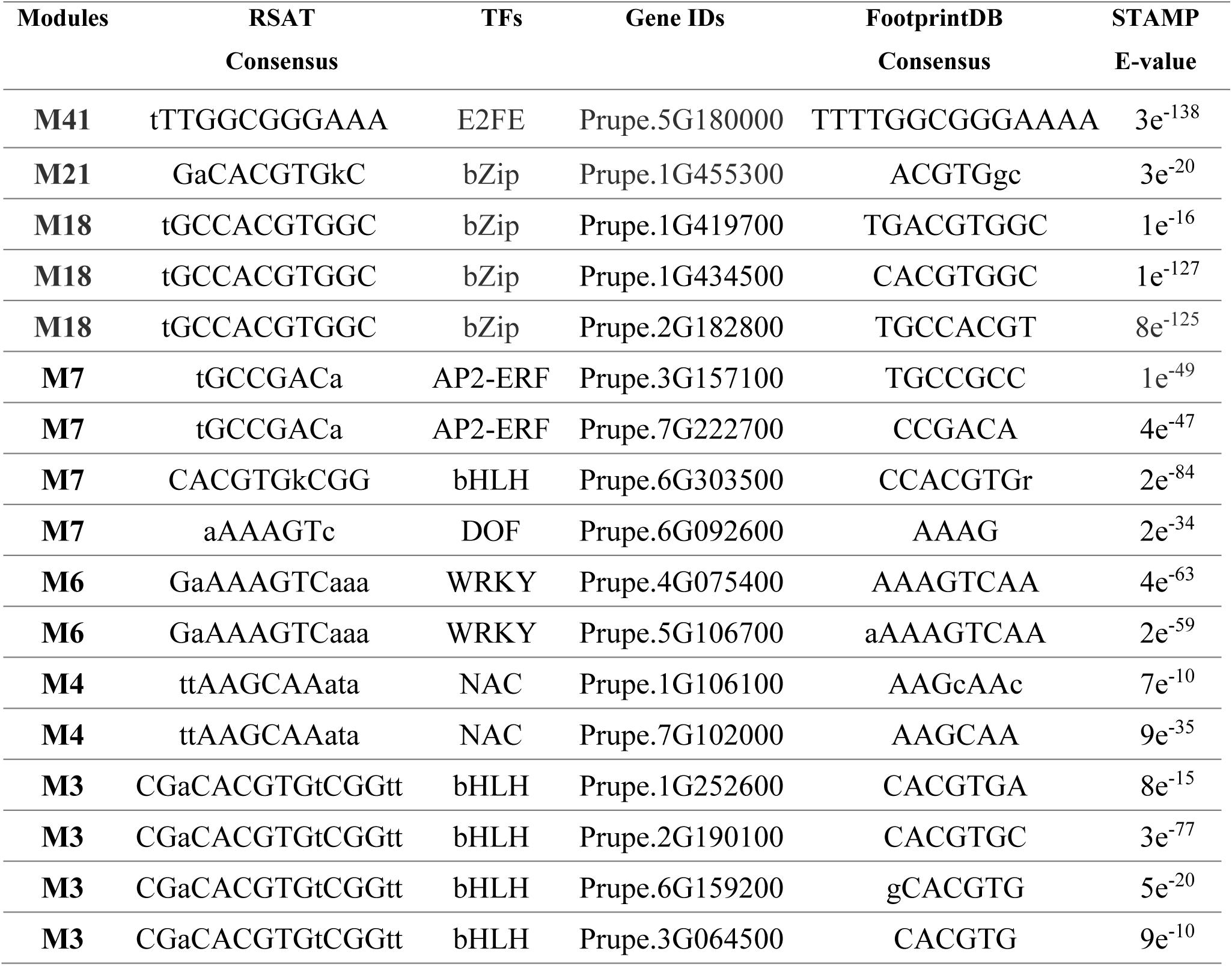
Similarity comparison between RSAT and footprintDB DNA-binding motif predictions. The best predictions in footprintDB were selected in *Arabidopsis thaliana*. The TFs grouped in this table are the same labeled with a star in **Figure 5**

### 2.5 Motif scanning

To identify the position of transcription factor binding sites (TFBS) in the promoter region of *P. persica* genes, position-specific scoring matrices (PSSMs) of all candidate motifs (77) were *in silico* scanned to the long (Up 1) upstream stretch [-1500, +200 bp]. We observed a clear positional bias of the TFBS close to the TSS, more precisely within the interval [-500 bp, +200 bp], then it progressively declines towards the 5’ limit (**Figure 6**). For motifs detected respectively in Up 1 (yellow color), Up 2 (green) and Up 3 (blue), sites were notably concentrated upstream the TSS showing a bell-shaped distribution from -500 bp to +0 bp with a maximum of density around -250 bp. Conversely, the positional distribution of motifs predicted along the upstream 4 was biased toward downstream the TSS with the flatter peak reaching its limit at the TSS (Up 4, purple). Detailed scanning results can be accessed at https://eead-csic-compbio.github.io/coexpression_motif_discovery/peach. On the other side, (AT) repetitive elements were also scanned to check their relevance, e.g., whether they correspond to the TATA box. The underlying data included in **Figure S2**, showed that TFBSs of these motifs were remarkably distant to the TSS and were distributed across the whole proximal region.

**Figure 6.**
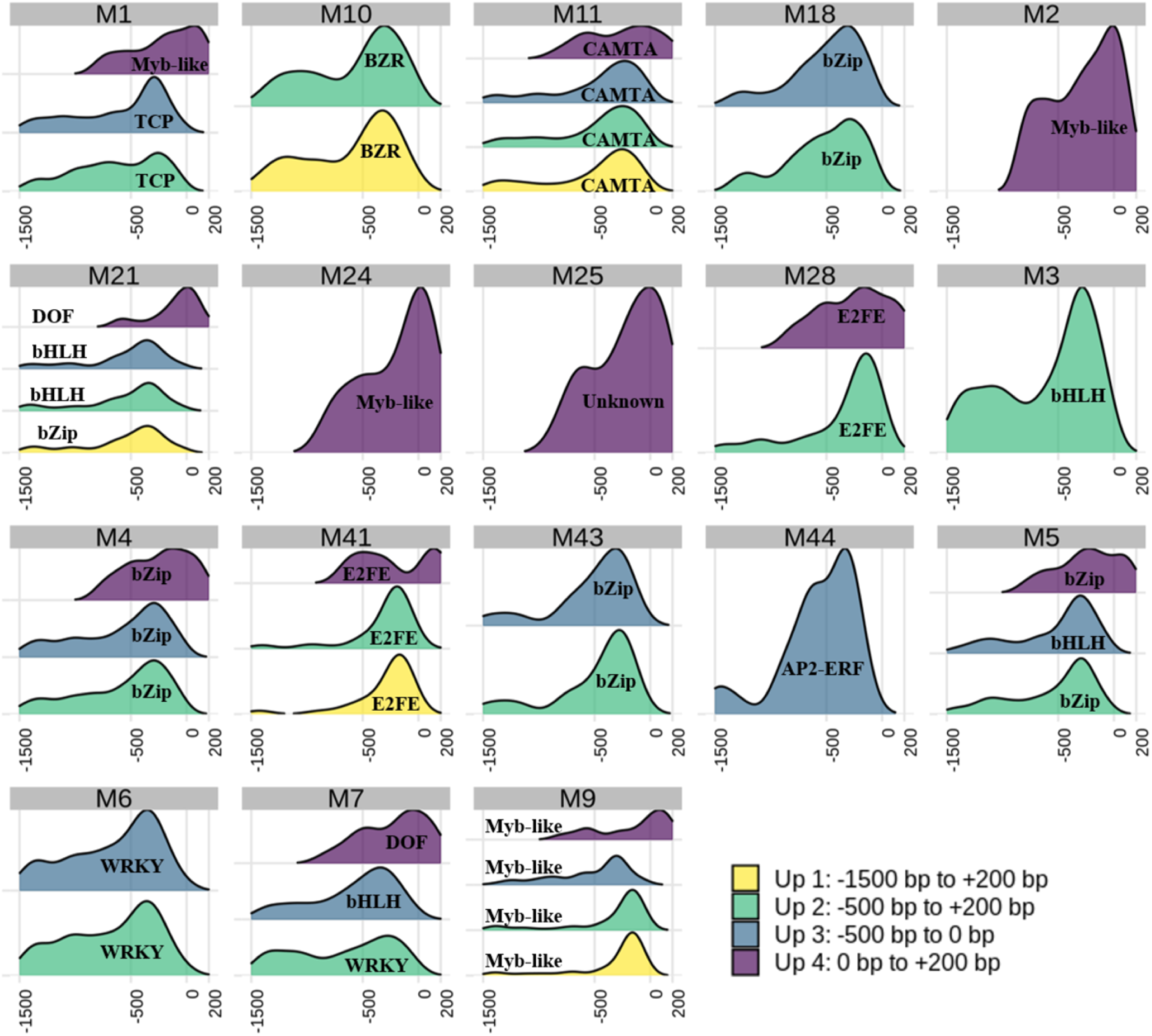
Positional distribution of the detected oligo motifs in promoter genes of *Prunus persica*. Four density distributions were derived from four assessed upstream regions. Up 1: from -1500 bp to 200 bp, Up 2: from -500 bp to +200 bp, Up 3: from -500 bp to 0 bp and Up 4 from 0 bp to + 200 bp. The x-axis corresponds to upstream length in base pairs (bp). The y-axis corresponds to density of captured sites with *P*-value <10 e^-4^. Only oligo motifs are presented here, dyads are provided in the report at https://eead-csic-compbio.github.io/coexpression_motif_discovery/peach.

### 2.6 Validation of the protocol for *de novo* cis-element discovery

To demonstrate the performance of the motif finding approach, we evaluated the effect of variable proximal promoter lengths on uncovering true DNA-binding sites in *Arabidopsis thaliana*. Experimentally proven motifs from a selection of *A. thaliana* transcription factors belonging to different families were successfully recovered by at least one algorithm. As summarized in **Figure 7**, JASPAR and *de novo* identified motifs displayed high consensus similarity. Moreover, in order to refine the comparison, we annotated the newly reported motifs JASPAR to ensure that they correspond to the TF family in question. As expected, *de novo* motifs shared the same annotation as the reference JASPAR motifs, which underlines the predictive performance of the proposed methodology.

**Figure 7.**
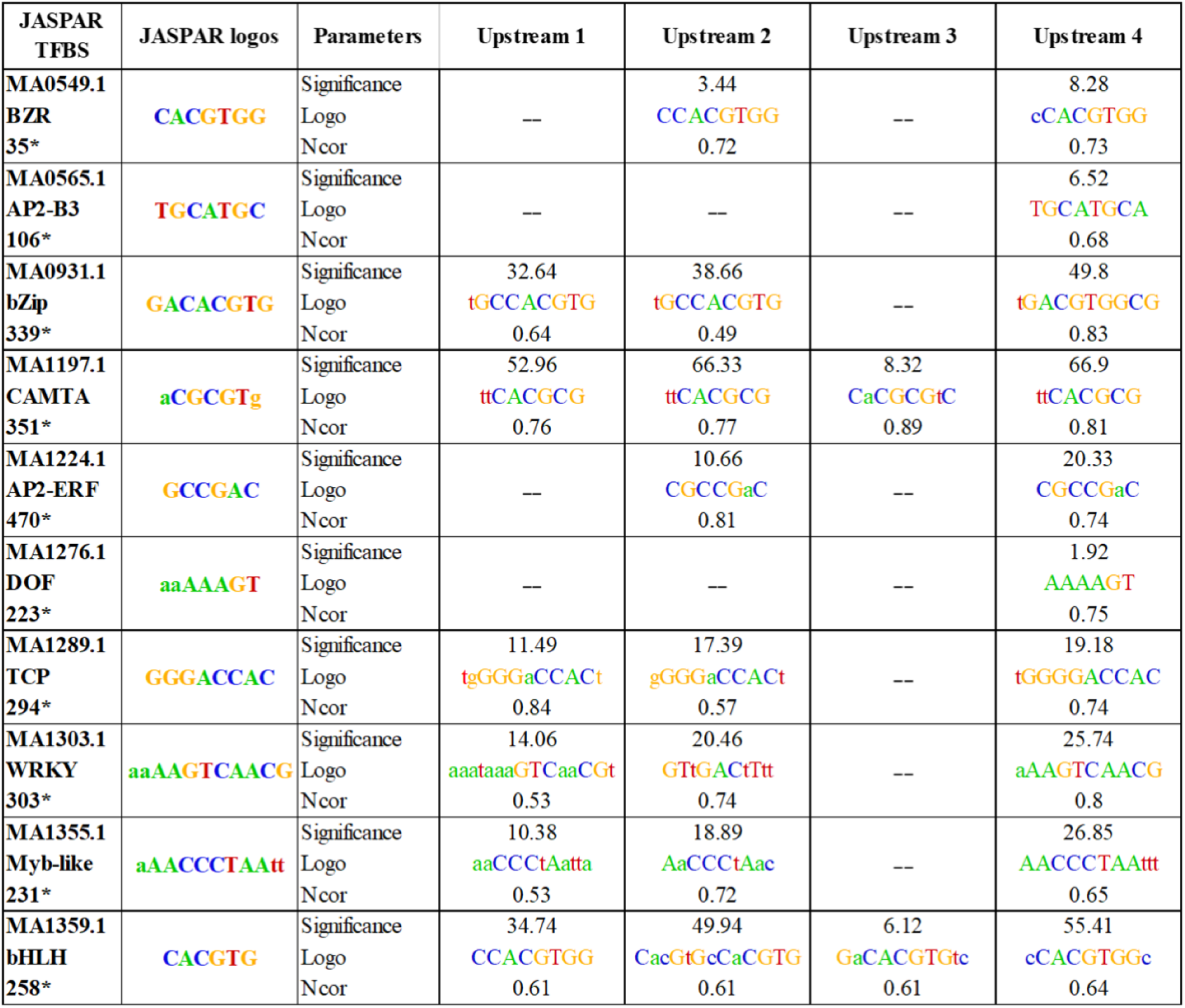
Similarity between JASPAR motifs (considered as queries) and *de novo* predicted oligo motifs found in *Arabidopsis thaliana* along four different upstream regions. Numbers tagged with a star indicate number of peaks recovered by BLASTN (see Methods). The Ncor scores correspond to JASPAR databases. Only oligo-analysis motifs are shown (dyads are available at supplementary **Table S4**). Upstream 1: [-1500 bp to +200 bp], Upstream 2: [-500 bp to +200 bp], Upstream3: [-500 bp to 0 bp] and Upstream4: [0 bp to +200 bp]

## 3. Discussion

In the present study, transcriptional profiling of eight independent data sets was conducted to decipher the intricate process of gene regulation in peach and to reveal meaningful biological signatures. DETs were grouped into 45 co-expression modules undergoing similar changes in their expression patterns. Unlike conventional clustering methods (such as k-means and hierarchical clustering), which are based on geometric distances, WGCNA is a graph-based approach relying on network topology as inferred from the correlation among expression values (Li et al., 2018). In our hands, the WGCNA algorithm robustly and accurately defined modules within a complex multi-condition dataset.

Discerning regulatory signals from blocks of co-expressed genes is a common presumption used to identify functional genomic elements. It has been successfully applied and approved in various plants species like *Arabidopsis thaliana* (Koschmann et al., 2012; Ma et al., 2013), *Zea mays* (Yu et al., 2015) and *Hordeum vulgare L*. (Cantalapiedra et al., 2017). However, little is known about its applicability to woody species. To our knowledge, this article is the first in which this hypothesis has been tested in *Prunus persica* genome wide.

For each predicted module, two-motif discovery algorithms (oligo and dyad analysis) were ran to discover significant motifs in the upstream promoter region. As suggested by Bianchi and colleagues, we initially defined the upstream promoter size as an interval of [-1500 bp to +200 bp] relative to the TSS (Bianchi et al., 2015). Discovered motifs with significant poly-(AT) sites were discarded due to their low complexity and scarcity of information concerning their specific-regulatory function. We reasoned that low complexity sequences might be linked to repetitive stretches of DNA, extensively present in plant genomes (Yu et al., 2015). Interestingly, when tuning the promoter upstream length to a tract of [-500 bp, +200 bp] relative to the TSS, these low complexity motifs were limited to module M2. It would seem that long upstream promoter regions unbalance the signal-to-noise ratio exacerbating the identification of such AT motifs. Along the same lines, we observe a dependence of (AT)-rich sites on the dataset size. Indeed, AT-low-complexity motifs were only detected in the first six modules, which contain from 560 to 1795 upstream sequences. In light of these considerations, we believe that in our study case, they may result in part due to the properties of DNA sequences (both upstream region length and dataset size) rather than the performance of the chosen algorithms. In **Table S3**, the results revealed that AT-rich occurrence in random cluster increases in parallel with the module size.

To check whether the AT-rich patterns overlap the TATA boxes, a position scanning experiment was conducted. It is well documented in plants that a TATA box region lays between -30 and +35 bp with respect to the TSS (Zhu Qun et al., 1995; Smale, 2001) However, the scanning results portrayed that peaks were located far from this interval, confirming that they are distinct signals (**Figure S2**).

By limiting the promoter length to a window of -500 bp, new regulatory motifs were recovered. Additionally, splitting the proximal promoter region into two intervals around the TSS enabled the discovery of further hidden candidate TF motifs. Such observations may strengthen our hypothesis that shorter upstream regions improve the sensitivity motif discovery (from 11 motif sequences identified within Up 1 to 58 sequences identified in Up 3 and Up 4 assessed separately). Defining the upstream promoter length has been a controversial issue (Kristiansson et al., 2009). If the interval is too short or too long, the motif of interest may not be captured. Therefore, we reason that an analysis on regions of variable length would yield a more comprehensive picture of the complex regulatory code.

The spatial distribution of the occurrences of the 77 inferred motifs along the promoter region is crucial to understand gene regulation in *Prunus persica*. Our findings revealed that TFBSs are not uniformly dispersed across the promoter but they exhibit a strikingly mixture of 2 density profiles: while the majority showed bell-shaped distribution at the interval of [-500 bp, 0 bp], others were diverged downstream the TSS [0 bp, +200 bp] (**Figure 6**). These findings are similar to those described in *A. thaliana*, with nearly two thirds of the examined TFBSs within the region from -400 bp to +200 bp (Yu et al., 2016). TFBSs of bHLH, BZR, TCP and WRKY are particularly concentrated from -500 bp to 0 bp. This denotes a positional binding preference within this proximal region, which is in agreement with (Yu et al., 2016) reporting that their positional preference is between -100 bp to -50 bp. On the other hand, bZip, CAMTA, E2FE and Myb-like exhibited a dual binding distribution with central peaks upstream and downstream the TSS. A possible explanation of this is that some TFs may display different binding preferences depending on their TF-specific structure, biological functions or combinatory with other TFs. The degree to which the arrangement of motif sites is associated to their function needs to be further investigated especially that data about TFBS distribution in plants is only limited to *Arabidopsis thaliana* (Zou et al., 2011; Yu et al., 2016). According to our findings, we may consider that the boundary from - 500 bp to 0 bp is an adequate region to look for the majority of TFBSs lying in the proximal promoter region in peach. However, we should keep in mind that proximal TFBSs could also occur downstream the TSS. Thus, we suggest defining the peach proximal promoter length as a tract of [-500 bp to +200 bp], analyzing separately the two regions around the TSS for a better motif coverage. In fact, according to Montardit-Tardà (2018), differences in the nucleotide composition were found upstream and downstream the TSS. At this point, we should mention that gene regulation involves a complex interplay between the proximal (promoter) and distal regulatory regions located thousands of base pairs away from the TSS (e.g., enhancers) (Li et al., 2019). Our workflow sheds light mainly on sequence signatures extracted from the proximal promoter. Thus, it might not be adequate to study distal genomic elements.

Furthermore, rather than barely returning a list of significant motifs, our methodology assigned them to different modules to help shape a clear overview of the peach regulation code. Overall, we were able to distinguish 18 modules harboring 77 motifs from 11 TF families: bHLH, bZip, BZR, CAMTA, DOF, E2FE, AP2-ERF, Myb-like, NAC, TCP and WRKY. While some modules, such as M6, M11, M28 and M41, seem to be driven by a single TF (WRKY, CAMTA and E2FE, respectively), motifs from different families were annotated in the rest. This can be explained by the fact that some promoter sequences may encompass multiple TFBSs of perhaps interacting TFs. Indeed, TFs have been reported to frequently operate in combination (Guo et al., 2018; Kumar et al., 2018). Combinatorial regulation is required to confer specific responses in a particular tissue and under a particular stress. Thus, the hypothesis of cooperative interactions between diverse motifs in peach is worthy to be further investigated.

From the inferred list of motifs (**Figure 2**), we found similar binding sequence potentially perceived by different class of transcription factors. For example, motifs “tGaCACGTGtc” and “GaCACGTGkCGg” in module M5 are distinct but can be aligned despite different nucleotide frequencies in some positions. We presume that TFs from related families may have similar DNA recognition sequences, as reported for instance by Franco-Zorrilla et al., 2014 for Myb and AP2 TFs.

The biological significance of modules with significant identified signals was determined by Gene Ontology analysis and TF annotation. The enriched modules reflected many biological functions involved in abiotic stress responses such as cold acclimation, response to stress, response to water and response to hormone (**Figure 4**). In this context, modules M1 enriched for “photosynthesis” contained candidate Myb and Myb-related factors. These findings are in line with (Baldoni et al., 2015) reporting that Myb TF family is known to regulate drought tolerance and the stomatal movements in plants. bHLH binding sites were mainly disclosed in modules M3, M5 and M7 (**Figure 5**). Associated TFs among those were abundant under various stress conditions proposing a multi-functional role of bHLH. According to Bianchi et al., (2015), bHLH factors play a central role in flavonoid biosynthesis and cold acclimation in peach. Similarly, bZip TFs were found in both M18 and M21 and their transcripts were mainly over-represented in all experiments. Our results are supported by previous studies reporting that bZip were induced by various environmental cues. Indeed it was revealed that they play a pivotal role in responses to cold stress in peach and enhance water use efficiency in almond-peach rootstocks (Hu et al., 2018). WRKYs putative motifs were restricted to M6 and were exclusively activated in leaf tissue under hyper-hydricity (HH) stress. It is well known that HH leads to morphological abnormalities, such as brittle leaves (Carrillo Bermejo et al., 2017). We speculate that WRKY factors may be implicated in morphological damages produced by HH. Module M11 was found to be a potential CAMTA-driven module (**Figure 2**), where two genes coding CAMTA were annotated (**Figure 5**). A previous study in *A. thaliana* demonstrated that cold stress increases the level of calcium sensed by CAMTA (Doherty et al., 2009). This perturbation of calcium levels leads to modification of the CAMTA activity that in turn triggers the induction of cold response genes of the CBF family. For this reason, CAMTA motifs are of great interest. From the perspective of peach breeding, these findings may be of great interest, as genes within modules are potential targets for further experimental validation.

Finally, a major drawback of motif discovery approaches is their limited performance. To tackle this issue we designed a control experiment in which genomic sites detected by ChIP-seq for 10 *A. thaliana* TFs were analyzed. Comparing the *de novo* predicted motifs to the corresponding curated motifs in JASPAR we observed a high similarity in terms of Ncor scores (**Figure 7 and Table S4**). When searching for *in-vivo* validated motifs, we would ideally expect to get identical predicted motifs. Nonetheless, while most consensus sequences had high Ncor values > 0.8, others had lower values. As well, we observed that the choice of upstream region length affects the performance. In some cases, particularly Up 1 and Up 3, the expected motif was not even found. Unlike the results found in peach, examining 4 upstream tracts only returned motifs from the same query families probably as a consequence of the JASPAR TFBSs profiles being curated. Taken together, we believe that the proposed workflow is robust enough to be extended to other species in order to identify reliable regulatory motifs.

## 4. Conclusion

DNA motif discovery is a primary step for studying gene regulation, however the *in silico* prediction of regulatory motifs in not straightforward. In contrast to previous surveys that usually assume a fixed promoter length right at the start; this work reports regulatory elements while testing different upstream sequence intervals. It is among the first efforts providing a comprehensive collection of *Prunus persica* motifs without a prior knowledge. By coupling gene expression networks and module analysis, we were able to extract interpretable information from a large set of noisy data and to reveal primary candidate TF-target binding sites responding to specific conditions. These results offer a more complete view of the proximal regulatory signatures in *P. persica* and we believe that it may contribute to address the knowledge gap about the transcriptional regulatory code in non-model species.

## 5. Materials and methods

### 5.1 Input data and processing

Eight peach RNA-sequencing datasets were downloaded from the European Nucleotide Archive (https://www.ebi.ac.uk/ena) and were used as raw reads for this project. This comprehensive dataset includes data of various peach cultivars, from various tissues (root, leaf, stigma and fruit), different stress conditions and developmental stages. A detailed list of the project IDs and metadata is provided respectively in **Table 1 and Table S1.A**. The obtained reads were quality-processed and trimmed using FASTQC v.0.11.5 and Trimmomatic v.0.36 (Bolger et al., 2014), to discard adaptors and low-quality sequences with mean Phred score (*Q* < 30) and window size of 4:15. The first nucleotides were then head-cropped to ensure a per-position A, C, G, T frequency near to 0.25. Following the trimming, only sequences longer than 36 bp were retained for further analysis. The complete workflow is shown in **Figure 1 (see step 1)**.

The high quality reads from each RNA-seq project were quantified separately using the pseudo-aligner kallisto v.0.43.1 for fast and accurate transcripts count and abundance (Bray et al., 2016). Kallisto was run in two steps: i) a transcriptome index was built from all cDNA transcripts of *Prunus persica* v2, from Ensembl Plants release 39 (Verde et al., 2017; Howe et al., 2020). ii) Each sample was pseudo-aligned against the index. Transcript level abundance was estimated and normalized to transcripts per million (TPM) using 100 bootstraps (-b 100) to ascertain the technical variation. For single-end read mode, average fragment length and standard deviation were additionally required and were set to (-l 200) and (-s 50), respectively.

### 5.2 Transcript-level profiling

Differential expression analysis was conducted with Sleuth R package v.0.29.0 (Pimentel et al., 2017) for each RNA data set separately. The Wald test (WT) was applied to output abundance files in order to retain the significant expressed transcripts from each experiment. Samples and their biological replicates from each experiment were compared with their corresponding control. To reduce the false positives, only transcripts passing an FDR cutoff *Q-*value < 0.01 and beta statistic (approximation of the Log2 Fold Change between two tested conditions) |β| > 1 were retained. Significant transcripts obtained from each RNA-seq project were merged into a single list with an assigned mean TPM value for each replicate.

### 5.3 Construction of co-expressed network

Based on the assumption that co-expressed genes may share the same biological signature, weighted gene co-expression network analysis (WGCNA v.1.61) was performed to extract clusters of densely interconnected genes named modules (Langfelder and Horvath, 2008). Samples were firstly clustered to remove outliers and transcripts with missing entries. A similarity matrix was constructed by performing pairwise Pearson correlation across all targets. Then an adjacency matrix was built raising the similarity matrix to a soft power (β). Here β was set to 7 reaching thus 83% of the scale free topology fitting index (R^2^). To minimize the effect of noise, matrix adjacency was transformed to Topological Overlap Measure (TOM) and its corresponding dissimilarity matrix (1-TOM) was generated. Finally, modules were defined using the cutreeDynamic function with a minimum module size of 20 targets. Compared to standard hierarchical clustering, this approach solves the issue of setting the final number of clusters and arranges the genes based on their topological overlap to eliminate spurious associations resulting from the correlation matrix.

### 5.4 *De novo* cis regulatory sequences discovery using RSAT::Plants

Gene modules resulting from network analysis were subjected to an *ab-initio* motif discovery pipeline using the RSAT::Plants standalone (**Figure 1, step 2**). For each module, the analysis initiates by generating as negative control 50 random clusters of the same size for each module as described previously (Contreras-Moreira et al., 2016). Sequences with four different boundaries around the TSS were retrieved from the genes in the co-expressed modules, random clusters and *Prunus persica* genome v2. The upstream sequences were defined as intervals of i) -1.5 kb to +200 bp ii) -500 bp to +200 bp and iii) two segments around the TSS: -500 bp to 0 and 0 bp to +200 bp. Note that the 0 to +200 interval corresponds to the 3’ UTR region, which is already downstream. RSAT *peak-motifs* was run under the differential analysis mode, where module’s upstream sequences served as the test set and all upstream sequences from peach genome were considered as the control set to estimate the background model (a background model was created for each upstream stretch) (Thomas-Chollier et al., 2012). Two discovery algorithms were used: i) oligo-analysis, which is based on the over-representation of k-mers in upstream regions, and ii) dyad-analysis, which looks for over-represented spaced pairs of oligonucleotides (Defrance et al., 2008). For each run, up to five motifs were returned per algorithm and were retained to compare their statistical significance with the 50 random clusters considered as negative control.

Candidate motifs were chosen based on their significance (log E-value) compared to negative control and were subsequently annotated by comparison to the footprintDB collection of plant curated motifs (http://floresta.eead.csic.es/footprintdb) (Sebastian and Contreras-Moreira, 2014) using the *compare-matrix* tool in RSAT (Nguyen et al., 2018) requiring a normalized correlation score Ncor ≥ 0.4.

Finally, selected motifs were scanned along the stretch [-1500 bp, +200 bp] to predict their corresponding binding site positions, using as background model a Markov chain of order 1 (m=1) and a cutoff *P*-value ≤ 1e^−4^. To ensure the clarity and reproducibility of this strategy, a repository including the source code, links to the results and a tutorial explaining how to reproduce a similar analysis on any species is available at https://eead-csic-compbio.github.io/coexpression_motif_discovery/peach.

### 5.5 Transcription factor prediction and Gene Ontology analysis

Hereafter, the analysis was restricted to modules with significant detected signals. Firstly, genes coding peach TFs were predicted and classified using the iTAK database (http://itak.feilab.net/cgi-bin/itak/index.cgi, last accessed January 2020). Protein sequences of TFs were subsequently submitted to footprintDB to predict their interacting DNA-binding site. To functionally interpret the co-expressed modules, Gene Ontology (GO) enrichment was conducted on PlantRegMap / PlantTFDB portal v5.0 (http://planttfdb.gao-lab.org/, last accessed January, 2020) (Tian et al., 2019). P-value of 0.01 was set to retain the significant GO terms.

### 5.6 Validation of the pipeline by detecting *a priori* known motifs in *Arabidopsis thaliana*

To assess the impact of upstream region lengths on the identification of relevant motifs, we used sets of experimentally validated binding sites of 10 *Arabidopsis thaliana* TF families. Sequences of the proven sites were downloaded from JASPAR database (Fornes et al., 2020) and were locally aligned with BLASTN against the *A. thaliana* TAIR10.42 genome from Ensembl Plants to obtain the closest neighbor genes. The following parameters were used: E-value ≤ 1e^-5^, max_target_seqs =1, max_hsps=1 query-coverage of 80% and percentage of identity 98%. Upstream sequences of neighbor genes were obtained with *retrieve-seq* from RSAT::Plants. Similarity between references (JASPAR) and newly discovered motifs was computed with Ncor score (see above).

## Supplemental Data

**Supplemental Table S1. A**. Detailed information about the RNA-seq data used for differential analysis

**Supplemental Table S1. B**. Number of survived and dropped reads after quality processing and pseudo-aligned reads using kallisto program

**Supplemental Table S2**. List of candidate regulatory sites discovered within four upstream tracts of different lengths. Motifs are represented as IUPAC consensus sequences. TF match: Transcription factor family of the best match in footprintDB. Ncor: normalized Pearson correlation varying between 0 and 1. Ncor ≥ 0.4 indicates high confidence annotations. Gray color indicates that no significant motifs were found. **Supplemental Table S3**. List of low complexity motifs considered as false positive predictions within a boundary from -1500 bp to +200 bp upstream region length. For each algorithm, sequences are presented both as IUPAC consensus sequences using the degeneracy code and as sequence logos. Last column indicates the occurrence number of AT-rich motifs within the 50 random clusters used as negative control. Ps: **W** letter refers to (A or T) nucleotide and **S** refers to (C or G). Number of sites corresponds to the occurrence number of a single motif.

**Supplemental Table S4**. Similarity of JASPAR motifs (considered as queries) and de *novo* predicted dyad motifs in *Arabidopsis thaliana*. Numbers tagged with asterisks indicate number of peaks recovered by BLASTN (see Methods). The Ncor scores correspond to JASPAR databases.

**Supplemental Figure S1**. Co-expression network analysis. **(A):** Sample clustering to detect outliers. Sample with the same node color are derived from the same RNA-experiment. **(B):** Topological overlap measure plot. The different shades of color signify the strength of the connections between the genes (from white not significantly correlated to red signifying highly significantly correlated). Modules identified are colored along both column and row and are boxed. **(C):** Distribution of the module size.

**Supplemental Figure S2**. Positional distribution of AT-rich repetitive motifs along upstream 1: [-1500 bp, +200 bp].

## Acknowledgements

We thank Eric OLO NDELA for his support and help in creating the HTML report and providing useful feedback. We also thank Claudio Antonio Meneses Araya, Dayan Sanhueza and Tomás Carrasco for giving us access to the RNA-seq data.

## Authors’ contribution

NK, BC-M and YG devised the study objectives, designed the experiment, discussed data and wrote the manuscript. NK performed the bioinformatics analysis, FM-T contributed to delimit the proximal promote region. JA-CM aided to prepare the figures and provided critical feedback. JvH contributed in the critical discussion of results. YG and BC-M contributed the analysis tools and YG conceived the experiment and supervised the activities. All authors read and approve the manuscript.

